# Optogenetic stimulation reveals frequency-dependent resonance and encoding in V1 excitatory and inhibitory neurons

**DOI:** 10.1101/2023.04.10.536138

**Authors:** Ana Clara Broggini, Irene Onorato, Athanasia Tzanou, Boris Sotomayor-Gómez, Cem Uran, Martin Vinck

**Affiliations:** Ernst-Strüngmann Institute for Neuroscience in Cooperation with Max Planck Society, Frankfurt Am Main, Germany; Donders Centre for Neuroscience, Department of Neuroinformatics, Radboud University Nijmegen, 6525 Nijmegen, the Netherlands

## Abstract

Cortical information processing is thought to be facilitated by the resonant properties of individual neurons and neuronal networks, which selectively amplify inputs at specific frequencies. We used optogenetics to test how different input frequencies are encoded by excitatory cells and parvalbumin-expressing (PV) interneurons in mouse V1. Spike phase-locking and power increased with frequency, reaching a broad peak around 80-100Hz. This effect was observed only for Chronos, a fast-kinetic opsin, but not for Channelrhodopsin-2. Surprisingly, neurons did not exhibit narrow-band resonance in specific frequency-ranges, and showed reliably phase-locking up to 140Hz. Strong phase-locking at high frequencies reflected non-linear input/output transformations, with neurons firing only in a narrow part of the cycle. By contrast, low-frequency inputs were encoded in a more continuous manner. Correspondingly, spectral coherence and firing rates showed little dependence on frequency and did not reflect transferred power. To investigate whether strong phase-locking facilitated the reliable encoding of inputs, we analyzed various spike-train distances and Fano factor. Interestingly, responses to lower rather than higher frequencies had more globally reliable spike-counts and timing structure. These findings have various practical implications for understanding the effects of optogenetic stimulation and choice of opsin. Furthermore, they show both PV and excitatory neurons respond with more local precision, i.e. phase-locking, to high-frequency inputs, but have more globally reliable responses to low-frequency inputs, suggesting differential coding regimes for these frequencies.

## Introduction

The cortex generates intrinsic oscillatory activity in distinct frequency bands that play an important role in temporally coordinating the activity of inhibitory and excitatory neurons (Buzsáki and Draguhn, 2004; Fries, 2015; Wang, 2010). Rhythmic activity in different frequency bands can be facilitated by the resonant properties of individual neurons and neuronal networks, which selectively amplify inputs at specific frequencies (Onorato et al., 2020; Gray and McCormick, 1996; Pike et al., 2000; Cardin et al., 2009; Lewis et al., 2021; Spyropoulos et al., 2022; Buzsaki and Voroslakos, 2023; Vinck et al., 2023). Resonance may play an important role in facilitating computation and selective information processing in cortical networks (Buzsaki and Voroslakos, 2023; Vinck et al., 2023; Izhikevich et al., 2003; Rusch and Mishra, 2020). Understanding the resonance properties of circuits in awake animals requires stimulation of neurons with cell-type specificity and at high temporal resolution. A promising tool to this end is optogenetics, whereby input currents into cells of specific classes can be generated with high temporal precision (Boyden et al., 2005; Emiliani et al., 2022). Thereby, neural transfer functions can be studied in a causal manner with cell-type-specificity, providing a deeper understanding of the input/output functions of cortical networks. This knowledge is not only crucial for advancing our understanding of the brain, but also for developing clinical applications of brain stimulation.

Previous studies using optogenetics have suggested that parvalbumin-expressing (PV) interneurons and excitatory neurons exhibit resonance specific frequency-ranges, most prominently the theta (4-12Hz) and gamma (30-80Hz) frequency-range. In a seminal study, Cardin et al. (2009) stimulated either excitatory or PV cells with optogenetics using ChR2 and quantified resonance at the network level using LFP power. Photoactivation of PV resulted in particularly strong LFP-power increases at gamma-frequencies, suggesting a pivotal role for PV interneurons in generating gamma oscillations. By contrast, photo-activation of excitatory neurons led to great power transfer at low frequencies. The latter observation agrees with other studies showing theta-resonance in excitatory neurons (Sirota et al., 2008; Voloh et al., 2015; Wang et al., 2006; Vaidya and Johnston, 2013; Stark et al., 2013). However, another recent study found gamma-resonance via optogenetic activation of excitatory neurons in V1 of anesthetized cats, as gauged from multi-unit activity (Lewis et al., 2021). Such resonance may be a particular feature of the visual cortex, which has the propensity to generate narrow-band oscillations (Onorato et al., 2020).

The previous work on optogenetics focused on the transferred power in response to optogenetic input stimuli of different frequencies, measured using LFPs or multi-unit activity (Lewis et al., 2021; Cardin et al., 2009) Yet, transferred power can be influenced by various factors, including: the elicited firing rate, e.g. more spikes elicited for higher frequencies; the synchronization of responses; and the linearity/non-linearity of neural responses. This raises the question, of how inputs at different frequencies are precisely encoded by excitatory and inhibitory neurons, and at which temporal precision. This question is important considering that network oscillations result from precise temporally synchronized firing among inhibitory and excitatory neurons (Gray and McCormick, 1996; Hutcheon and Yarom, 2000; Cardin et al., 2005; Buzsáki and Wang, 2012).

To investigate these questions, we performed high-density recordings in V1 of the awake mouse, while photo-activating neurons at different frequencies with sinusoidal or white-noise stimuli. We used an opsin with fast kinetics, Chronos, and compared the responses with ChR2. Furthermore, we compared the effects of photo-activating excitatory and PV neurons. Well-isolated single neurons were identified based on short-latency responses to the optogenetic stimuli. For these neurons, spike phase-locking and power increased with frequency, reaching a broad peak around 80-100Hz. However, neurons did not exhibit narrow-band resonance in specific frequency-ranges. Strong phase-locking at high frequencies reflected non-linear input/output transformations, with neurons firing only in a narrow part of the cycle. Correspondingly, spectral coherence and firing rates were very weakly modulated by input frequency and did not reflect transferred power. Finally, to investigate whether strong phase-locking facilitated the reliable encoding of inputs, we analyzed various spike-train distances and Fano factor. We show that responses to lower rather than higher frequencies had more globally reliable spike-counts and timing structure.

## Results

We used high-density silicon probes (Cambridge Neurotech, see Methods) to record spikes and Local Field Potentials (LFPs) from area V1 in awake mice. Mice were placed on a running wheel and were passively viewing a grey screen (Figure 1a). Recordings were performed in four different groups of mice, which expressed one of two different opsins (Chronos or ChR2) in either excitatory (using AAV-CaMKII) or PV+ neurons (parvalbumin+; in PV-Cre mice). During the recordings, we applied light stimulation via an optical fiber placed on the brain surface. For each mouse, we used four different stimulation protocols, namely DC (direct current), white noise, pulses and sinusoidal stimuli of different frequencies (Figure 1a). We then analyzed the responses of well-isolated single units (sorted semi-automatically via Kilosort) to the different light stimuli. It is known that V1 activity shows a strong dependence on behavioral state (McGinley et al., 2015; Niell and Stryker, 2010).

**Figure 1:**
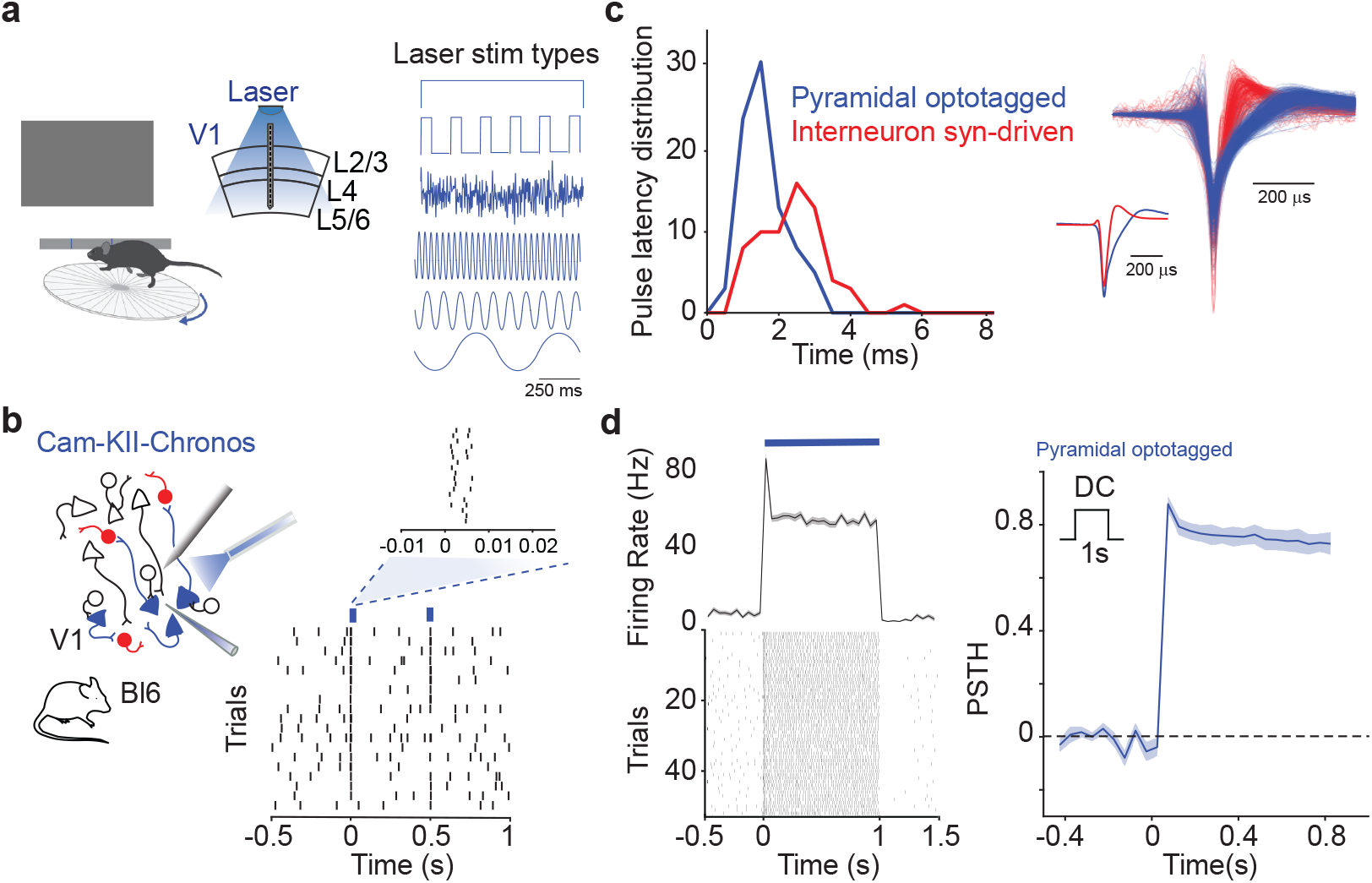
Optotagging of excitatory neurons that expressed the fast opsin Chronos. **a**) Illustration of the experimental paradigm. Awake mice were placed on a running wheel, and passively viewing a grey screen. Single neurons and LFPs were recorded from area V1 using a high-density silicon probe. In a subset of time epochs, laser stimuli were applied through an optical fiber. Four different types of photoactivation protocols were used: DC (direct current), white noise, short pulses, and sinusoidal stimuli of different frequencies. **b**) Excitatory neurons (labeled in blue on the illustration on the left) expressed the Chronos opsin under the regulation of the CaMKII promoter. The raster plot shows the response of chronos-expressing neurons to optogenetic pulses and based on the short-latency response observed (note on the top right inset), these neurons were identified as optotagged cells (N=87 cells). **c**) Distribution of response latencies to short pulses (5ms) stimuli. The neurons that were identified as Chronos-expressing (i.e. Pyramidal optotagged neurons) had broad-waveform action potentials and fired at an earlier latency (1.6ms) after light onset than fast-spiking (2.4ms, Two-sample t-test, p = 7.8 × 10^-7^), narrow-waveform (NW) interneurons. The NW interneurons are called “Interneurons syn-driven” because of the indirect activation mediated by excitatory neurons. **d**) Example raster plot and average PSTH of optotagged single neurons showing a sustained increase in firing rates during the presentation of DC light stimuli (1s).

This raises the question of whether the behavioral state of the animal, running or static, would affect the neuronal resonance at different frequencies and other aspects of the input/output function. However, we did not observe differences in the firing rates and phase locking between running and static periods (Figure S1). Hence, for the main analyses, we pooled behavioral states together.

### Optotagging

In mice with the expression of Chronos in excitatory neurons, we identified two main sets of neurons: First, so-called “optotagged” neurons (N = 87 cells) that had broad-waveform (BW) action potentials and short-latency responses to optogenetic pulses, which indicated that these neurons were directly photo-activated (Figure 1b-c, see Methods). Second, so-called “syn-driven” fast-spiking (FS) interneurons (N = 73 cells) that had narrow-waveform (NW) action potentials and which increased their firing during the presentation of DC stimuli. These FS interneurons fired with a short latency (2.4ms) of about 1 ms after the optotagged neurons (1.6ms), indicating that these FS interneurons were not directly photo-activated, but mono-synaptically activated by the light-driven excitatory neurons (p<0.01, Figure 1c-d).

Optotagged neurons showed a sustained increase in firing during the presentation of DC stimuli (Figure 1d). This result differs substantially from a previous study that analyzed multi-rather than single-unit activity, and concluded that light-driven responses mediated by Chronos are highly transient (Jun and Cardin, 2020).

### Neurons phase-lock most precisely to high-frequency inputs

Next, we analyzed firing responses to sinusoidal stimuli of different frequencies (Figure 2a). These sinusoidal stimuli had equal light power at all frequencies (see Methods). To analyze the phase-locking of spikes to the sinusoidal stimuli, we first determined the phase of each spike relative to the laser cycle (i.e., using the equality *φ* = 2*πft*). The strength of phase-locking was then quantified with the pairwise phase consistency (PPC; (Vinck et al., 2012)), which is a method that is unbiased for the number of spikes. In addition, we computed the spike train’s output power, using the Discrete Fourier Transform, at the stimulation frequency plus the higher harmonics.

**Figure 2:**
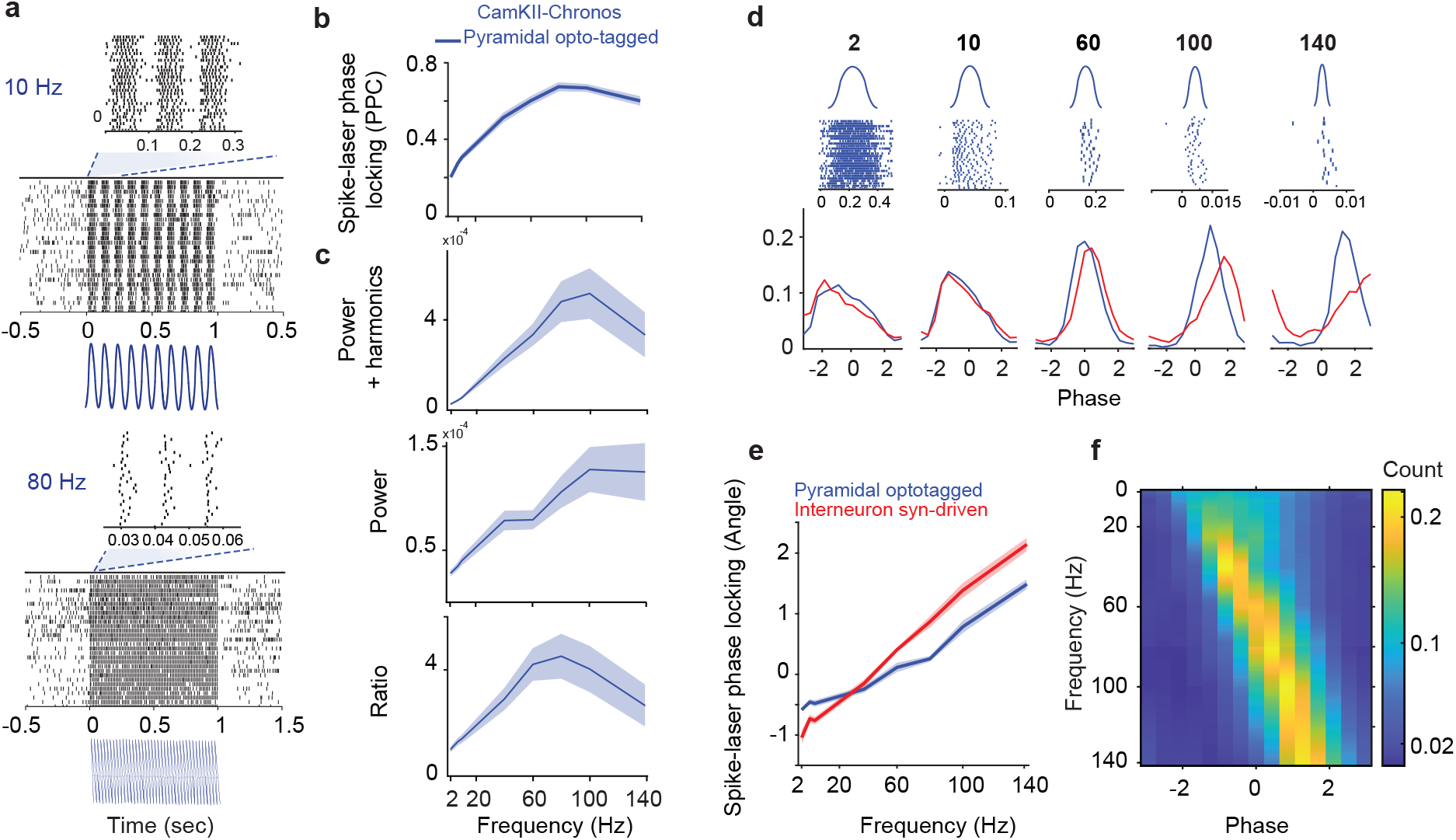
Phase-locking increased towards higher frequencies, without band-pass gamma resonance. **a**) Example neuron with its firing responses to Chronos sinusoidal stimuli of 1s duration at 10 and 80 Hz. Insets zooms in on 3 cycles at 10 and 80 Hz. **b**) Spike laser phase-locking increases towards higher frequencies. Average of Spearman correlation coefficients across cells between PPC values and frequencies = 0.71, p<0.01. Spike-laser mean PPC at 80Hz differed significantly to PPC at 140Hz (t-test, p=0.004). **c**) Similar increase observed for the spike train’s output power at the stimulation frequency (Power), plus the higher harmonics (Power+harmonics) and for the ratio of these 2 measures. Average of Spearman correlation coefficients across cells between Power values and frequencies = 0.44, p<0.01, between Power+harmonics and frequencies = 0.46, p<0.01. **d**) Distribution of spike phases relative to the laser cycle. Top, example neuron’s raster plots at frequencies 2, 10, 60, 100 and 140Hz. Bottom, average of phase distribution across cells for pyramidal optotagged cells (blue trace) and interneurons synaptically driven (red trace). **e**) Mean of spike-laser phase locking angle at the laser stimulation frequencies for pyramidal optotagged cells and interneurons synaptically driven. **f**) Mean of spike phases relative to the laser cycle for pyramidal optotagged cells (73 cells) from frequency 2-140Hz.

Across neurons, phase locking increased strongly with frequency and reached a plateau around 80-100 Hz (Figure 2b). From 100 to 140 Hz, phase locking only showed a minor decrease (Figure 2b). The spike train’s power spectrum showed a similar profile (Figure 2c).

To further understand the mechanisms underlying the increase in phase-locking with frequency, we further characterized the distribution of spike phases relative to the laser cycle. We observed that phase distributions became increasingly narrow for higher frequencies, consistent with the corresponding increase in phase-locking (Figure 2d-f). By contrast, at low frequencies, optotagged neurons showed a substantially wider phase distribution (Figure 2d), consistent with the relatively weak PPC values (Figure 2b). Hence, the increase in phaselocking can be attributed to the fact that at high frequencies, neurons are active only in a very narrow part of the light cycle, whereas at low frequencies, they track the sinusoidal light stimulus in a more continuous i.e. linear manner.

Inspection of the phase distributions further revealed a nonlinear relationship between the laser input and the firing activity of the optotagged neurons. At low frequencies, the optotagged neurons fired with a phase advance relative to the laser stimulus (Figure 2d-f). At high frequencies, optotagged neurons fired relatively late in the laser cycle. At high frequencies, we further observed a clear phase delay between the optotagged neurons and the FS interneurons, which fired about 1 ms later than the optotagged neurons, consistent with a mono-synaptic activation of the FS interneurons.

We further examined how the dependence of phase-locking on frequency related to the changes in firing rates. To determine this, we computed the average firing rates during the light period and compared them to the baseline period. Firing rates increased only weakly across frequencies, and this increase was only significant when including frequencies below 20Hz (Figure 3a). This finding suggests that the increased phase-locking is not due to an increased drive of neurons at higher frequencies, but rather due to enhanced precision.

**Figure 3:**
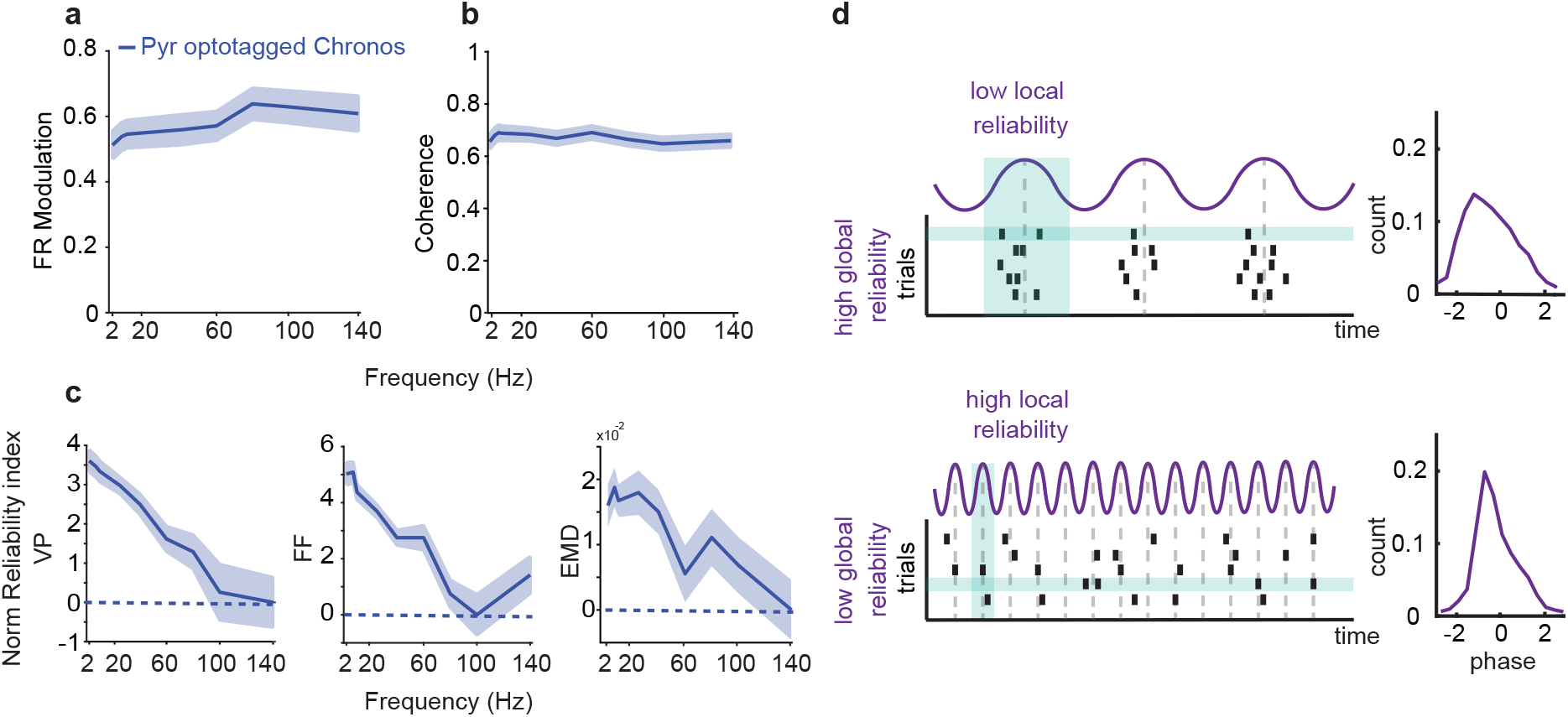
Different properties of neuronal resonance: firing rate (FR) modulation, spectral coherence and neuronal reliability. **a**) Firing rate (FR) modulation of optotagged neurons by sinusoidal photoactivation using Chronos at different frequencies. The FR modulation was computed as (*a* - *b*)/(*a* + *b*). Here *a* is the FR average during laser stimulation and *b* the FR average during baseline (pre-laser stimuli period). Spearman’s correlation coefficient, rho=0.18, p= 0.02. However this correlation is driven by inclusion of low frequencies, e.g. above 20Hz correlation is n.s. (rho = 0.073, p = 0.36). **b**) Spectral coherence between the spike train and the laser at different frequencies. Spearman’s correlation coefficient rho= −0.22, p= 0.004. **c**) Measurements of neuronal reliability estimated as normalized indices. Three different types of reliability were computed: Victor Purpura distance (VP, left plot), Fano factor (FF, middle plot) and Earth mover distance (EMD, right plot). The VP distance is a generic spike train distance incorporating both rate and timing difference. Fano Factor measures spike count variability normalized by average spike count. The Earth Mover Distance computes the minimum transport cost between spike trains strictly based on the timing structure. These measurements were normalized by subtracting from the value of reliability per cell, the average across frequencies. Then, the indices were computed as (*mm* - *mf*). Here *mm* is the maximum value of reliability across frequencies and *mf*, the mean value or reliability per each frequency. Spearman’s correlation coefficient for VP, FF, EMD, rho= −0.98, rho = −0.95, rho = −0.73, respectively. p<0.001 for the three correlation measurements. **d**) Scheme representing two different regimes of reliability. Local and global reliability, with a high degree of local (i.e. precise spike phase locking) reliability for higher frequencies, and high global (i.e. reproducable spike trains) reliability for low frequencies

Finally, we wondered how the phase-locking compared to the spectral coherence between the spike train and the laser. Coherence reflects the extent to which one signal can linearly predict another signal, whereas phase locking reflects the width of the spike-phase distribution. We found that the spectral coherence showed a minor but significant decrease across frequencies (Figure 3b). Considering that coherence is a measure of linear prediction, this result indicates that the optotagged neurons contained slightly more information about the stimulus input in their spiking output at low than high frequencies. That is the enhanced phase locking at high frequencies, resulting from a narrow phase distribution, did not translate into an improved linear encoding of the laser input.

### Analysis of responses to white-noise stimulation

The analysis of responses to sinusoidal stimuli reveals a broad peak in the phase-locking spectrum, reaching a plateau around 80-100Hz. However, we did not observe narrow-band resonance in the gamma-frequency range. We wondered whether a similar conclusion would be obtained using white-noise stimulation, as in Lewis et al. (2021). White-noise photo-activation (low-pass filtered at 150Hz) resulted in sustained increases in spike rates in opto-tagged neurons (Figure S3A). However, the coherence spectrum between spike trains and white noise stimuli was approximately flat, consistent with the absence of resonance (Figure S3B). Analysis of the spike-triggered average showed that spikes were preceded by rapid depolarizations in the light stimulus with a short delay (2.44 ms). The spike-triggered average did not show rhythmic modulation that differed from the short-lived ringing imposed by low-pass filtering of the white noise (Figure S3C). These results are consistent with the finding that for sinusoidal stimuli, there is no narrow-band resonance. Note that for white-noise stimuli we did not perform an analysis of phase-locking, because the instantaneous phase cannot be defined for white noise.

### Reliability of neuronal responses

We observed that neurons showed a strong increase in phaselocking for higher frequencies. At first glance, this suggests that neurons respond very reliably to fast temporal fluctuations in the input, at least in terms of the phase at which the neurons fire. However, the precision of firing as a function of phase does not necessarily imply that neurons also show reliable responses to the inputs *across repetitions* of the same input. Notably, it was reported by Mainen and Sejnowski (1995) that in slices, neurons show very similar responses to repeated presentations of the same input pattern when it contained higher-frequency stochastic fluctuations. We wondered if the optotagged neurons show a similar degree of reliability to the same input pattern across repetitions.

To quantify this, we computed three different measures. First, the Victor-Purpura (VP) distance, which is a generic spike train distance measure that reflects the minimum cost of transforming one spike train into another spike train (see Methods), and covers both insertion and shift costs (Victor and Purpura, 1996,1997). Furthermore, we computed the Fano Factor which measures reliability only based on firing rates (i.e. variance / rate) (Eden and Kramer, 2010; Charles et al., 2018; Rajdl et al., 2020) and obtained a normalized measure of reliability (see Methods). Last, we computed the Earth mover distance (EMD), which is a way to measure the distance between probability distributions by calculating the minimum cost needed to move mass from one distribution to another (Grossberger et al., 2018; Sotomayor-Gómez et al., 2023). EMD can be used to measure the similarity or dissimilarity of spike trains by treating them as probability distributions. In this context, the spike trains are represented as point processes, and each spike is considered a unit of mass. The three measures were computed between all pairs of spike trains across trials (i.e. each pair of trials *k* and *j*) and then averaged and normalized them to obtain a measure of reliability. We found that neuronal reliability showed a monotonic decrease with frequency, given that all the 3 measures analyzed presented a strong negative correlation with the frequencies (Figure 3c; for PV neurons see Figure S2). Thus, although neurons had a high “local” reliability for high-frequency inputs in terms of the consistency of spike phases, the neurons had a low “global” reliability for high-frequency inputs in the sense that the spike trains across repeated presentations were very different from each other (Figure 3d).

### Stimulation of excitatory neurons using ChR2

The results described above were obtained with the opsin Chronos, which has different properties including faster kinetics. We thus wondered how the responses of neurons to different stimuli depended on the choice of the opsin. For example, the widely used opsin ChR2, which was used in previous studies examining frequency-dependent responses to optogenetic stimuli (Lewis et al., 2021; Cardin et al., 2009), has slower kinetics compared to the recently introduced opsin Chronos.

We performed similar experiments as for Chronos and identified a sample of optotagged excitatory neurons. We then examined the response of these neurons to the DC stimuli, as a previous study has suggested that in area V1, ChR2 yields more sustained responses than Chronos (Jun and Cardin, 2020). In contrast to the study by Jun and Cardin (2020), we found that photo-activation with DC light stimuli yielded sustained responses (Figure 4b) that were comparable to the experiments with Chronos.

**Figure 4:**
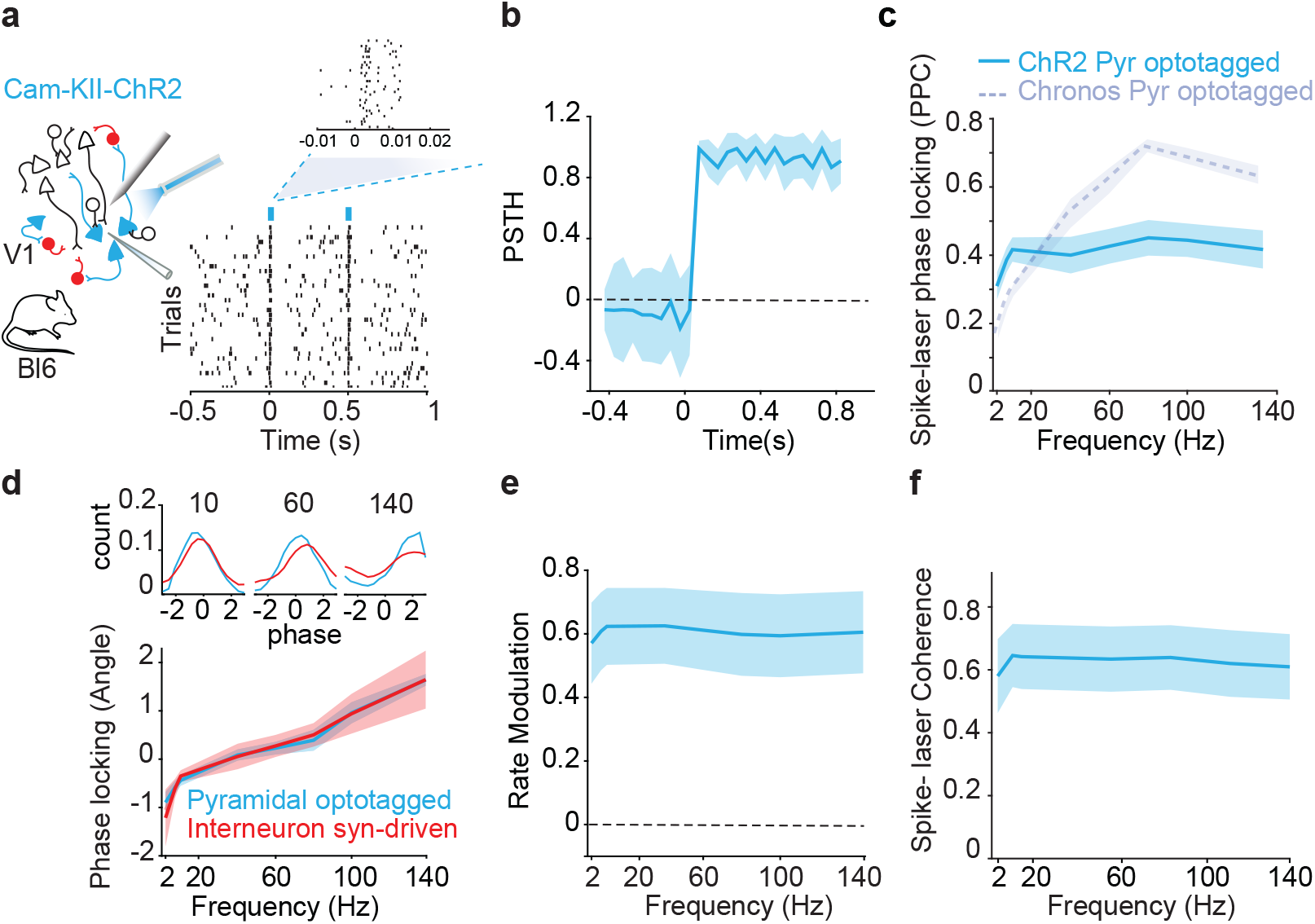
Resonance properties depend on the opsin used during optogenetics stimuli. **a**) Excitatory neurons expressed the Chr2 opsin under the regulation of the CaMKII promoter. Chr2-expressing neurons (N=17 cells) were identified based on short latency responses to optogenetic pulses (see Methods). **b**) ChR2-Optotagged single neurons showed a sustained increase in firing rates during the presentation of DC light stimuli. **c**) Flat spike-laser phase locking spectrum across frequencies when stimulating excitatory cells expressing ChR2 (light blue solid line). Average of Spearman correlation coefficients across cells between PPC values and frequencies = 0.3782, p=0.03. Spike-laser mean PPC at 80Hz did not differ from PPC at 140Hz (t-test, p=0.99). Chronos expressing neurons spike-laser phase locking was plotted again here for comparison (grey dashed line, Spearmam’s rho coefficient= 0.71, p = 0.004). **d**) Top, Average of phase distribution across cells for pyramidal optotagged cells (blue trace) and interneurons synaptically driven (red trace). Bottom, Mean of spike-laser phase locking angle at the laser stimulation frequencies for pyramidal (Pyr) optotagged cells(blue trace) and interneurons synaptically driven (red trace). **e**) FR modulation of ChR2-optotagged neurons by laser sinusoidal stimuli at different frequencies. Neurons were equally modulated by different types of frequencies. Spearman’s correlation coefficient, rho=0.4152, p= 0.002. FR modulation at 80Hz did not differ to FR modulation at 140Hz (t-test, p=0.43) **f**) Spectral coherence between the spike train and the laser at different frequencies. Spearman’s correlation coefficient rho= −0.26, p= 0.15.

Next, we predicted that because of its kinetics, ChR2 should cause additional low-pass filtering compared to Chronos. Indeed, we found that the phase-locking of spikes to the laser did not show an increase with frequency, and was substantially weaker at higher frequencies (Figure 4c). Thus, the preferred phase-locking to higher frequencies only became apparent when using an opsin having fast kinetics (Figure 4c).

Finally, we found that several other aspects of the neural responses, i.e. firing rates, coherence, and reliability, were similar comparing Chronos and ChR2 (Figure 4c-d, Figure S2).

### Stimulation of PV+ neurons

Finally, we wondered how neural responses to sinusoidal stimuli differ between cell classes, in particular excitatory neurons and PV+ (Parvalbumin+) interneurons. Indeed, a previous study in area S1 of anesthetized mice using the opsin ChR2 has suggested that there is network resonance in the gamma-frequency range when photo-activating PV+ interneurons, but not when photo-activating excitatory neurons (Cardin et al., 2009).

We, therefore, repeated the experiments in two other sets of mice using either the opsin ChR2 or Chronos expressed in PV+ interneurons (Figure 5a-b). Surprisingly, we found that PV+ interneurons were substantially less phase-locked to the sinusoidal light stimuli than excitatory neurons, both for Chronos and ChR2 (Figure 5c). In addition, we did not observe an increase in LFP power (Figure 5d), specifically for photo-activation at gamma frequencies, in contrast to Cardin et al. (2009). Thus, we conclude that there is no specific resonance in the gamma-frequency range when stimulating PV+ interneurons as compared to excitatory neurons, neither at the level of individual neurons nor at the level of the network.

**Figure 5:**
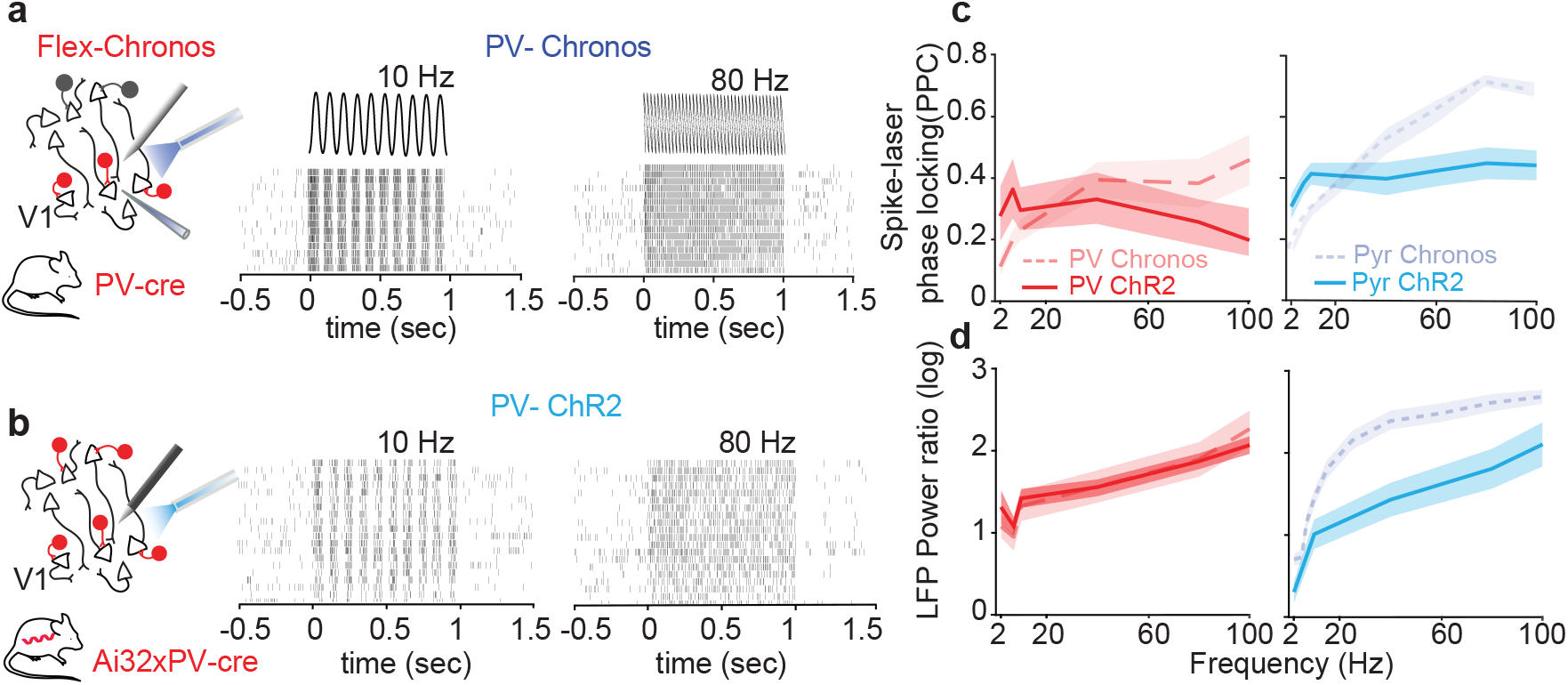
PV+ interneurons were substantially less phase locked to the sinusoidal light stimuli than excitatory neurons. **a**) PV+ cells expressed the Chronos opsin after injection of Flex-Chronos virus to PV-Cre mice, allowing Cre-dependent expression of the opsin (see Methods). Raster plot of example neuron showing PV+ cells firing responses to Chronos sinusoidal stimuli of 1s duration at 10 and 80 Hz. **b**) The same as panel **a**, but here PV+ cells expressed the opsin ChR2. The crossing of the genetic lines Ai32xPV-cre mice were used for Cre-dependent mediated expression of ChR2. **c**) Spike-laser phase locking spectrum across frequencies when stimulating PV+ cells expressing ChR2 (red solid line) or Chronos (light red dashed line). Average of Spearman correlation coefficients across cells between PPC values and frequencies was not significant for ChR2-Pv+ cells (rho = −0.15, p=0.7). Similarly, the mean spike-laser PPC observed at 60Hz or 80Hz did not differ significantly from that observed at 100Hz (t-tests, p=0.86 and p=0.45, respectively). Average of Spearman correlation coefficients across cells between PPC values and frequencies for Chronos-Pv+ cells, rho = 0.73, p<0.001. The mean spike-laser PPC observed at 60Hz or 80Hz did not differ significantly, also for Chronos-Pv+ cells, from that observed at 100 Hz (t-tests, p=0.38 and p=0.98, respectively). Spike-laser phase locking of Chronos and ChR2 excitatory neurons were plotted again here for comparison (on the right, pyramidal (Pyr) cells expressing Chronos are represented by solid blue line, ChR2 by light blue dashed line). Note that excitatory neurons are more locked to phase locked to the sinusoidal light stimuli than PV+ cells. **d**) LFP power ratio (LFP power during laser stimulation/ LFP power during baseline) for PV+ cells expressing ChR2 (red solid line) or Chronos (light red dashed line). Average of Spearman correlation coefficients across cells between LFP power ratio values and frequencies was significant for PV+ Chronos and PV+ Chr2 cells, rho = 0.82, p<0.001 and rho = 0.83, p<0.001, respectively. LFP power ratio (LFP power during laser stimulation/ LFP power during baseline) for pyr cells expressing ChR2 (blue solid line) or Chronos (light blue dashed line). Average of Spearman correlation coefficients across cells between LFP power ratio values and frequencies was significant for pyr Chronos and pyr Chr2 cells, rho=0.93, p=0 and rho = 0.86, p<0.001, respectively.

## Discussion

We investigated how neurons in mouse V1 respond to optogenetically-induced inputs at different frequencies, using an opsin with fast kinetics. We found that both excitatory and inhibitory neurons show the strongest phase locking at frequencies of 80Hz and above, which translated into an increase in spike train power. We did not find evidence for band-limited resonance in the gamma-frequency range, neither for excitatory nor inhibitory neurons. The transformation of inputs to outputs showed two kinds of non-linearities: On the one hand, a phase advance and asymmetric phase distribution at lower frequencies, and on the other hand, a narrowing of the phase distribution for higher frequencies corresponding to an increased transfer to higher harmonics. In contrast to phase locking, we found that firing rates were largely stable across frequencies, suggesting a form of normalization whereby neurons are equally well driven by all frequencies. Furthermore, increased phase locking at higher frequencies did not result in a higher coherence, but slightly lower coherence, indicating that spike trains provided equal linear predictions for all frequencies. While neurons fired with high local reliability (i.e. a reliable phase distribution) to high frequencies, we found that neuronal firing became increasingly “globally” unreliable at higher frequencies. Here global reliability was quantified in terms of Fano factor and spike train distance (Victor-purpura distance and spike timing distance). Thus, there is a tradeoff between local and global reliability, with a high degree of local (i.e. precise spike phase locking) reliability for higher frequencies, and high global (i.e. reproducible spike trains) reliability for low frequencies. The latter finding suggests that neural populations may utilize different coding strategies for lower and higher frequencies.

### Reliability

Previous work has suggested that *in vitro*, stimulation with repeated high-frequency patterns causes highly reliable activation patterns (Bryant and Segundo, 1976; Mainen and Sejnowski, 1995). However, our results indicate this situation is markedly different *in vivo*. We observed that firing responses were generally more reliable across repetitions for low-rather than high-frequency stimuli. This held true for reliability quantified in terms of firing rates (Fano factor), and spike train distance measures that partially (VP-distance) or entirely (EMD) reflect the temporal structure of spike trains. Thus, high-frequency stimuli are effective in driving neurons and causing local phase locking, however, yield highly variable response patterns in terms of the average spike count and the global timing within the spike train. Hence, it appears to be a trade-off between local reliability (in terms of phase locking) and global reliability (in terms of the general temporal structure of the spike train).

This finding may suggest that neural populations use different mechanisms to encode inputs at lower and higher frequencies. At lower frequencies, individual neurons show reliable responses and encode inputs in a more gradual manner. As the phase-locking on a local scale (i.e. within a given cycle) is relatively weak, the transferred power appears weak. However, the stimulus is encoded with relatively high fidelity and can be read out in individual cycles. At higher frequencies, individual neurons phase-lock very strongly which results in high signal power. The increased phase-locking reflects a narrowing of phase distributions, rather than an increase in firing rates, and neurons encode the input in a highly non-linear manner. In this case, individual neurons show a high degree of stochasticity in the responses across cycles and trials, such that the encoding of a given cycle must rely on the response of a population of neurons. Our observations fit well with the idea that at high frequencies, neural populations are essentially in a noise-driven regime allowing for high entropy and stochasticity (Buzsaki and Voroslakos, 2023).

### Input-output functions characterized using optogenetics

Optogenetics provides a valuable tool for the identification of input-output functions in individual neurons and neuronal networks (Lewis et al., 2021; Cardin et al., 2009). Yet, the nature and extent of perturbations that can be induced are limited by the biophysical characteristics of the employed opsins, such as their conductance, kinetics, and sensitivity (Jun and Cardin, 2020; Hass and Glickfeld, 2016; Yu et al., 2020). It should be noted of course that the filtering properties of the opsin are not flat, but do show some degree of low-pass filtering, esp. for opsins with slower kinetics. Furthermore, using optogenetics may bypass to some extent synaptic filtering properties that may impose additional low-pass filtering. There are various ways in which these input-output functions can be characterized. Here, we compared several aspects of the input-output function, including firing rate, phase locking, power, coherence, and reliability.

The classic measure of resonance is the transfer function, or the transferred power from the input to the output system, i.e. the gain. In our experiment, as the power was kept constant across frequencies, the gain can be measured through the power spectrum of the spike trains. We observed that the spike train power generally increased towards higher frequencies (i.e. 100Hz), both at the input frequencies as well as the higher harmonics (which indicates a non-linearity that we discuss further below). The power of the spike trains as a function of frequencies is the product of spike count (squared) and the phase locking, i.e. *P*(*f*) ∝ *n*(*f*)^2^ø(*f*).^2^, where *φ*(*f*) is the resultant vector length and n(f) is the spike count. That is, power at a given frequency can be increased due to two factors.

We separated these two factors. First, we found that, surprisingly, the firing rate showed a very weak dependence on the frequency of stimulation. This result indicates that the increase in power is not due to an increase in the spike count. The likely explanation is that at higher frequencies, periods of strong depolarization are more frequent, however, the cells are unable to fire at all of these cycles. We propose that these results may have interesting implications for computation, as they indicate that a similar amount of energy is devoted to represent inputs at different frequencies, which could correspond to a maximum entropy that maximizes information transmission.

Second, by contrast, the phase locking showed a very similar profile as the power spectrum, that is, it increased monotonically with increasing frequency. This indicates that the increase in power is explained by phase locking. Notably, we did not observe a narrow-band gamma-resonance, but rather an increase in phase locking that reached a plateau around 80-100Hz, and decreased only slightly until 140Hz. Our experiments with white noise stimuli also did not provide evidence for gamma-resonance. Thus, these findings suggest that the transfer function is characterized by a high-pass filtering profile, rather than a resonance at gamma frequencies. Such high-pass filtering is likely explained by the active integration properties of neurons, i.e. the spike-generating mechanism (Izhikevich et al., 2003). Our data indicate that this high-pass filtering has a non-linear component as reflected by narrow phase distribution and transfer of input energy to higher frequencies.

### Comparison with previous studies on gamma-resonance

These conclusions on gamma-resonance strongly differ from two previous studies by Cardin et al. (2009) and Lewis et al. (2021). Lewis et al. (2021) measured MUA and LFP activity in anesthetized cats and provided sinusoidal and white noise stimuli with expression of the ChR2 opsin. With sinusoidal stimuli, they observed that the modulation depth of the MUA autocorrelogram (Ni et al., 2017) and MUA-laser cross-correlogram (Lewis et al., 2021) reached a plateau at 40Hz and did not observe a decrease towards 80Hz. However, with white noise stimuli, they conclude that there is resonance in the gamma-frequency band. This conclusion is based on Granger-causality analyses and the determination of the transfer function. One potential explanation is that cat V1 contains a resonant circuit, driven by the presence of gamma-resonant pyramidal units (Onorato et al., 2020; Gray and McCormick, 1996). Another potential explanation, however, is that in the cat cortex, white noise stimuli elicit a strong gamma-band response which may be explained by the DC component. However, it is not excluded that the statistics adopted can be influenced by the presence of gamma in the MUA. In particular, the transfer function 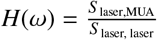 can be directly influenced by the magnitude of the MUA power spectrum. Likewise, Granger-causality can be influenced by the signal-to-noise ratio of the MUA which will peak in the gamma-frequency range. Thus, it remains unclear whether the data of Lewis et al. (2021) is indeed explained by genuine resonance to the white noise stimuli or rather reflects the induction of gamma by the DC component of the white noise stimuli.

Cardin et al. (2009) measured the transfer function at the level of the local LFP while stimulating PV+ interneurons in area S1 of the anesthetized mouse expressing ChR2. Cardin et al. (2009) reported that gamma-frequency stimulation caused a marked increase in LFP power, whereas lower and higher frequencies caused a smaller increase. Furthermore, stimulating excitatory neurons was accompanied by low-pass filtering. Similar to this study, we also stimulated PV+ interneurons either with ChR2 or Chronos in area V1 of the awake mouse, and we also quantified the change in LFP power. In contrast to Cardin et al. (2009), we did not find any evidence for gamma resonance at the level of spike trains or the LFP, but rather found evidence for high pass filtering. Interestingly, the input/output function of PV+ neurons that we observed was not markedly different from the input/output function of excitatory neurons. In fact, excitatory neurons showed stronger phase locking at higher frequencies than PV+ interneurons, which differs strongly from the findings of Cardin et al. (2009). It is unknown whether these differences are explained by the brain area recorded from (S1 vs. V1) or the behavioral state of the animal (anesthetized vs. awake).

### Comparison between opsins

An important factor in the study of resonant properties with optogenetics is the choice of the opsin. To our knowledge, the current study is the second one that compares neural response properties between ChR2 and Chronos *in vivo*, which are known to have slower and faster kinetics, respectively (Klapoetke et al., 2014). Jun and Cardin (2020) compared multi-unit responses in awake mice between ChR2 and Chronos. They reported that responses to DC stimuli were relatively sustained for ChR2, however, showed substantial adaptation for Chronos. Our results differ from this previous study, as we found that sustained responses could be generated also with Chronos. Hence, Chronos is suited for experiments in which sustained activation is required. A possible explanation for this difference is that in this we focused on the response properties of opto-tagged, well-isolated single neurons, whereas Jun and Cardin (2020) analyzed multi-unit responses. Another major difference that we described is that sinusoidal activation of neurons mediated by Chronos results in strong phase locking at high frequencies. By contrast, phase locking at high frequencies is substantially diminished in the case of ChR2, resulting in a flat phase-locking spectrum across frequencies. This reduction with ChR2 fits with the slower kinetics of this opsin. We also observed robust phase differences between excitatory neurons and putative fast-spiking interneurons for Chronos at higher frequencies, but not with ChR2 (Figure 4).

### Outlook

In sum, we do not find evidence for narrow-band theta or gamma resonance. Neurons respond with more local precision, i.e. phase-locking, to high-frequency stimuli, but show more reliable responses to low-frequency inputs, suggesting differential coding regimes for these frequencies. Our results may have several consequences for the use of optogenetics for manipulating circuits and for clinical purposes. First, our findings show that it is actually possible to entrain or sync a population at very high frequencies, even pyramidal neurons, by using an opsin with fast kinetics. A recent study has shown that such high-frequency stimulation may effectively ameliorate behavioral symptoms in Parkinsonian rats (Yu et al., 2020). Second, at high frequencies, the results of stimulation on a single neuron level are quite variable, which may predict that there is more variability in terms of behavioral effects. Third, high-frequency stimulation may have interesting consequences in terms of putting the circuit in a stochastic regime (i.e. have high variability) even though there’s strong phase-locking locally. Although we did not explore this, forcing the network in such a stochastic regime may also have specific consequences for synaptic plasticity. Fourth, our results suggest that low-frequency stimulation would be most effective to transmit information in a reliable manner, which could be important for e.g. BCIs.

## EXPERIMENTAL MODEL AND SUBJECT DETAILS

### METHOD DETAILS

#### Animals and virus injections

Experiments were performed on three to eight months old mice, both genders were used. All procedures complied with the European Communities Council Directive 2010/63/EC and the German Law for Protection of Animals and were approved by local authorities, following appropriate ethics review. Mice were socially housed with their litter on an inverted 12/12 h light cycle and recordings were performed during their dark (awake) cycle. The mice were mostly in a quiet, awake behavioral state. We expressed either Chronos (AAV1.CamKIIa.Chronos-eYFP-WPRE, VectorBiolabs) or ChannelRhodopsin-2 (ChR2, AAV1.CamKIIa.hChR2(H134R)-eYFP.WPRE.hGH, VectorBiolabs) under the CamKII promoter with the purpose of activating excitatory V1 neurons and also to identify these neurons through optotagging. For inhibitory neurons, we injected a Flex-Chronos virus (AAV1-Ef1a-Flex-rc[Chronos-GFP],VectorBiolabs) to PV-Cre mice (B6.129P2-PV+tm1(cre)Arbr/J, JAX Stock 017320, The Jackson Labaratory) to express Chronos in PV+. In order to compare the entrainment effects when modulating PV+ cells using a different opsin, we crossed the genetic mouse line Ai32(RCL-ChR2(H134R)/EYFP), which contains a CRE-dependent ChR2, with PV-Cre mice to allow Cre-dependent expression of ChR2 in PV+ (PV-ChR2).

#### Electrophysiological recordings

Thirty minutes prior to the head-post surgery antibiotic (Enrofloxacin, 10 mg/kg, sc, Bayer, Leverkusen, Germany) and analgesic (Metamizole, 200 mg/kg, sc) were administered. For the anesthesia induction, mice were placed in an induction chamber and briefly exposed to isoflurane (3% in oxygen, CP-Pharma, Burgdorf, Germany). Shortly after the anesthesia induction, the mice were fixated in a stereotaxic frame (David Kopf Instruments, Tujunga, California, USA) and the anesthesia was adjusted to 0.8 – 1.5% in oxygen. To prevent corneal damage the eyes were covered with eye ointment (Bepanthen, Bayer, Leverkusen, Germany) during the procedure. A custom-made titanium head fixation bar was secured with dental cement (Super-Bond C & B, Sun Medical, Shiga, Japan) exactly above the bregma suture, while the area of the recording craniotomy (V1, AP: 1.2 mm anterior to the anterior border of the transverse sinus, ML: 2.1 to 2.5 mm) was covered with cyanoacrylate glue (Insta-Cure, Bob Smith Industries Inc, Atascadero, CA USA). Four to six days after the surgery, the animals were habituated for at least five days in the experimental conditions. On the same day of the first recording session a 0.6 mm2 craniotomy was performed above V1 (AP: 1.2 mm anterior to the anterior border of the transverse sinus, ML: 2.1 to 2.5 mm, coordinates obtained from Wang et al. (2011) and adjusted experimentally), under isoflurane anesthesia. The craniotomy was covered with silicon (Kwik-Cast, World Precision Instruments, Sarasota, USA), and the mouse was allowed to recover for at least 2 hours. Recording sessions were carried out daily for a maximum of 5 days, depending on the quality of the electrophysiological signal. Awake mice were head-fixed and placed on the radial wheel apparatus. For the electrophysiological recordings, we used both Cambridge Neurotech probes (optotagged cells = 87 cells). During the experiments, we recorded simultaneously from 64 channels, in order to record from all the layers of V1 (up to around 800 μm). During the optogenetic experiments, an optic fiber (Thorlabs, 200μm, 0.39 NA) coupled to a diode laser (LuxX CW, 473 nm, 100 mW, Omicron-Laserage Laser produkte GmbH, Germany) was used to illuminate V1 craniotomy. The optic fiber was positioned 0.2 mm from the probe position, just above the surface of the brain.

#### Behavioral state quantification

We analyzed locomotion trials that had an average speed of more than 0.6 cm/s until the next point where the locomotion stopped, and also lasted for at least 2 seconds. Additionally, we selected quiescence trials that lasted for more than 5 seconds, had an average speed lower than 0.6 cm/s, and had a maximum movement range of less than 3 cm throughout the entire quiescence trial (criteria adapted from Vinck et al. (2015)).

#### Optogenetic stimuli and Optotagging protocol

The optogenetic manipulation consisted of 4 types of stimuli: short square pulses (5ms), sinusoidal (1s), continuous square light pulses (direct current, DC, 1s) and white noise. Short pulses and DC were applied in order to optotag cells. To investigate the entrainment of the network at different frequencies (2, 10, 25, 60, 100, and 140 Hz) we used sinusoid laser stimuli, ensuring equal power at all frequencies. DC stimuli were also used to estimate the intensity curve responses and choose the amplitude power applied for each recording session. For that, continuous light square pulses were applied for 1 second interleaved by 3-6 s intervals. The light intensity on the tip of the fiber was 3-80 mW/mm2, adjusted for each session according to neuronal responses. White noise stimulation was used to control for the presence of frequency-specific resonance effects. The optotagging protocol to tag pyramidal cells consisted of approximately 50 trials of 1s DC stimuli and 50 trials of 5ms square pulses stimulation periods with randomized inter-stimulus intervals between 4 and 6 seconds. Pyramidal optotagged cells were identified based on 3 different criteria: (1) Presence of a broad waveform, (2) a significant increase of firing probability after short square pulses (5ms) within 3ms after the light onset, and (3) positive firing rate modulation compared to baseline during the first 200ms of a continuous square light pulse (1s). The firing modulation was computed as FR(laser)-FR(pre-laser stimuli period)/FR(laser)+FR(pre-laser stimuli period) (Figure 1b-d). For criteria 2, the average of spike count should be statistically higher than random permuted matrix. The optotagging protocol to tag PV+ cells consisted of approximately 50 trials of 1s long DC stimulation periods with randomized inter-stimulus intervals between 4 and 6 seconds. PV+ optotagged cells were considered light-responsive if were fulfilling 2 criteria: (1) positive firing rate modulation compared to baseline during DC stimuli and (2) a significant increase of firing probability for at least 5ms bin, computed over the period of laser stimulation (excluding the first 5ms to avoid the presence of light artifacts) (Figure 5a-b).

#### Data analysis

##### Waveform classification

The mean waveform was calculated over data segments from −41 to 42 samples around the time of the spike, based on the aligned waveforms of the 10000 spikes randomly selected for each neuron. The sampling rate was increased by a factor of 3 using spline interpolation. The mean waveforms were normalized by subtracting the median of the first 10 samples and then dividing by the absolute value of the negative peak. Waveforms with a positive absolute peak were discarded. Subsequently, two-dimensional t-Stochastic Neighbor Embedding (t-SNE; perplexity of 80) was applied on the 80 samples after the negative spike peak of the waveforms. Lastly, we applied hierarchical clustering on the two-dimensional t-SNE embedding, which resulted in two separate clusters corresponding to the broad and narrow waveform neurons.

##### LFP preprocessing and Spectral analysis

To obtain LFPs, electrode signals were first low-pass filtered at 400 Hz and then high-pass filtered at 0.1 Hz, using a third-order Butterworth filter. In order to filter out line noise, an additional band-stop filter between 49.5 and 50.5 Hz and 99 and 101 Hz was applied. Subsequently, signals were downsampled to 1200 Hz by averaging consecutive frames.

##### Spike-laser phase locking

Single units (700 cells) were isolated using the semi-automated spike sorting algorithm Kilosort 2.5 (Steinmetz et al., 2021) and classified as putative excitatory and inhibitory neurons. We analyzed the phase-locking of single units to the laser using the PPC (pairwise phase consistency), a measure that is unbiased for the number of spikes. The PPC was calculated using windows of 250 ms around each spike (Vinck et al., 2012), using the ft spiketriggeredspectrum functions in the FieldTrip SPIKE toolbox.

##### Neuronal reliability

Three different measurements of neuronal reliability were computed: Victor Purpura distance (VP) (Victor and Purpura, 1996, 1997), Fano factor (FF) (Eden and Kramer, 2010; Charles et al., 2018; Rajdl et al., 2020) and Earth mover distance (EMD) (Grossberger et al., 2018; Sotomayor-Gomez et al., 2023). These measurements were normalized by subtracting from the value of reliability per cell, the average across frequencies. Then, the indices were computed as (mm-mf), where mm is the maximum value of reliability across frequencies, and mf, is the mean value or reliability per each frequency.

##### Statistical Testing

Statistical details, including the specific statistical tests and p values, are specified in the corresponding figure legends or results section. In general, paired t-test, Wilcoxon-Mann-Whitney, and Wilcoxon signed-rank tests were performed. Throughout the whole paper, data are presented as the mean ±SEM, unless otherwise indicated. All statistical analyses were conducted using MATLAB 2020a (Mathworks).

## Authorship contributions

Conceptualization and design: AB, IO, MV. Analysis: AB, BS, MV. Recordings: AB, IO, AT. Surgery: AB, IO, AT. Materials and reagents: AT, CU, MV. Supervision: MV. Funding acquisition: MV. Writing initial draft: AB, MV. Review/Editing: All other authors.

## Declaration of Interests

The authors declare no competing interests.

---

**Figure S1:**
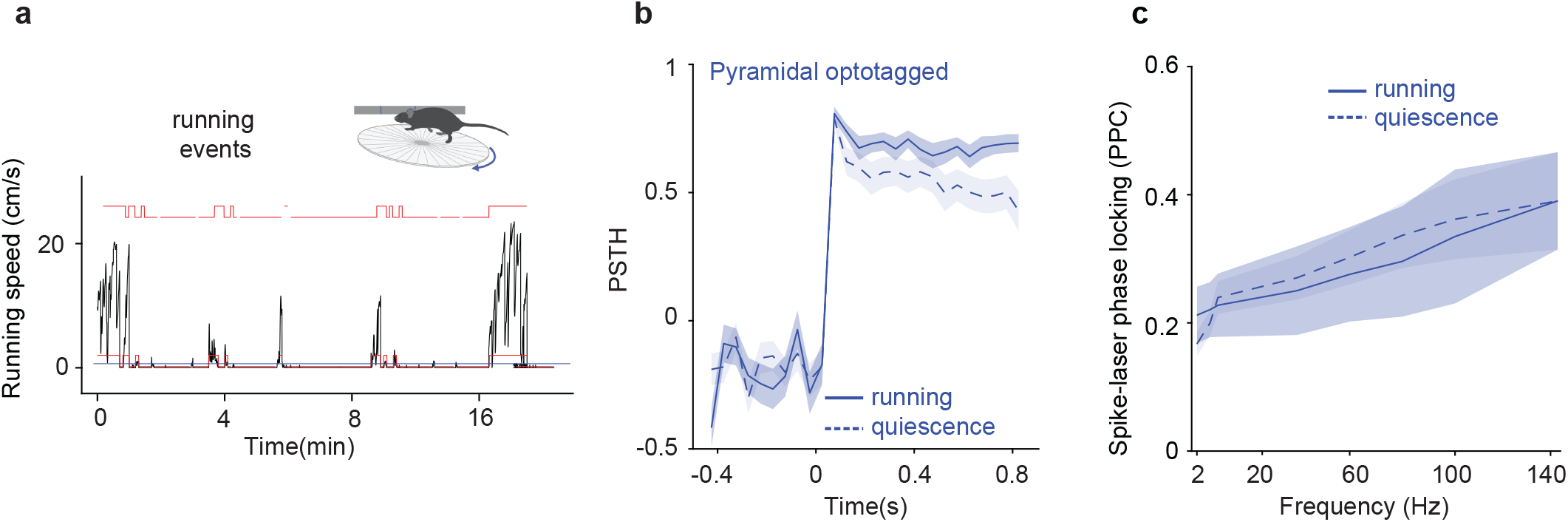
Average neuronal response and spike-laser phase locking during laser stimuli of Pyr cells expressing Chronos do not differ between running and quiescent states. **a**) Mouse running speed (cm/s) of an example session. The running speed of each trial period (1s) was estimated based on the change of radians read from the running wheel encoder while the animal was head-fixed seating/running in the running wheel. The estimated speed was then concatenated and the running events (red line) were defined as trials where the average velocity was higher than 0.6cm/s (plotted as the blue line in the figure). **b**) Average neuronal response to 1s squared pulse laser stimuli using Chronos opsin. The solid blue line shows the average response during running trials and the dashed blue line during quiescent trials. **c**) Mean of spike-laser phase locking at the laser stimulation frequencies for pyramidal optotagged cells during running trials (blue solid line) and quiescence trials (blue dashed line).

**Figure S2:**
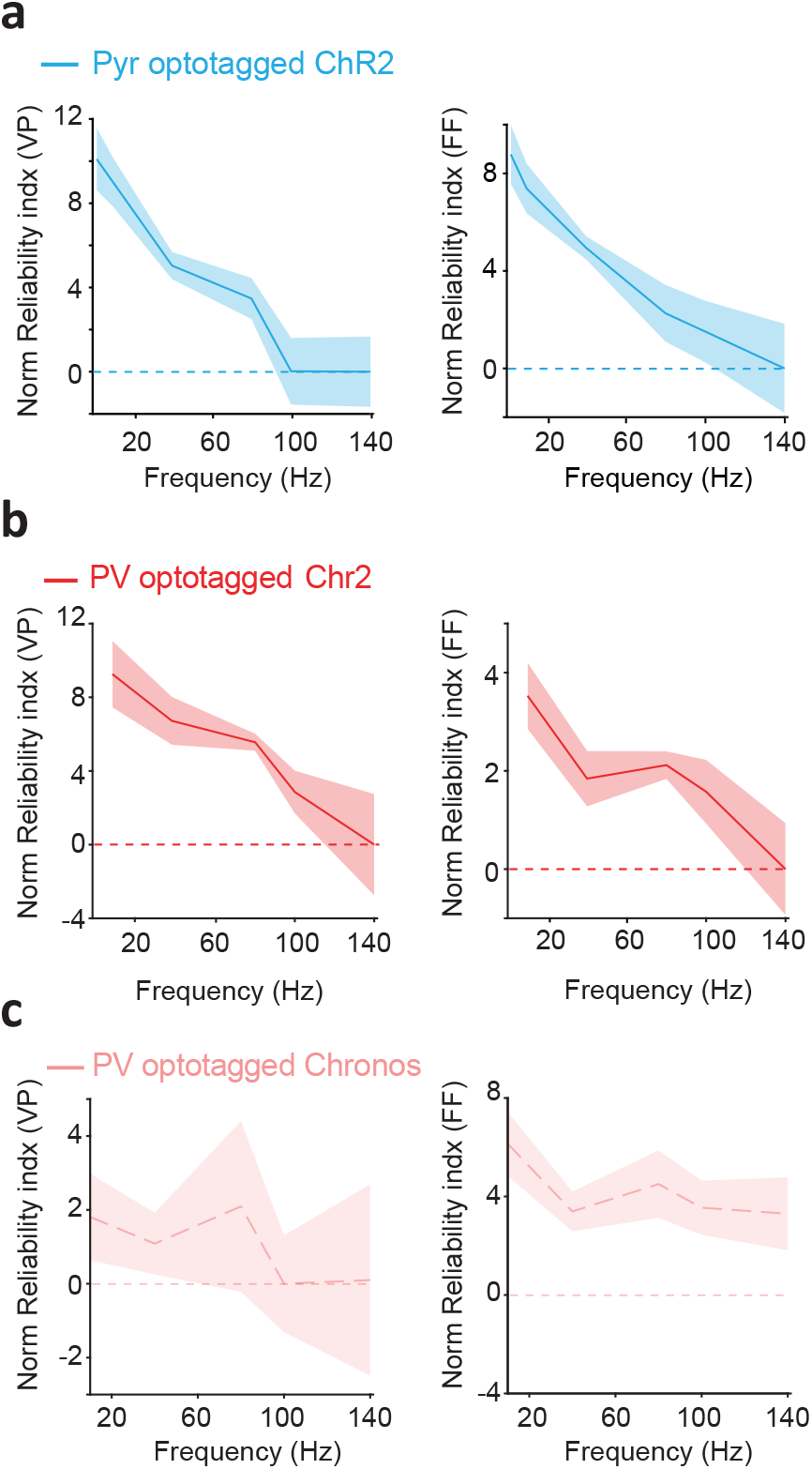
Measurements of neuronal reliability estimated as normalized indices. Two different types of reliability were computed: Victor Purpura distance (VP, left plot) and Fano factor (FF, right plot). These measurements were normalized by subtracting from the value of reliability per cell, the average across frequencies. Then, the indices were computed as mm-mf. Here mm is the maximum value of reliability across frequencies and mf, is the mean value or reliability per each frequency. **a**) Neuronal reliability of Pyramidal cells expressing Chr2 when optogenetically stimulated at different frequencies. **b**) Neuronal reliability of PV+ cells expressing Chr2 when optogenetically stimulated at different frequencies. **c**) Neuronal reliability of PV+ cells expressing Chronos when optogenetically stimulated at different frequencies.

**Figure S3:**
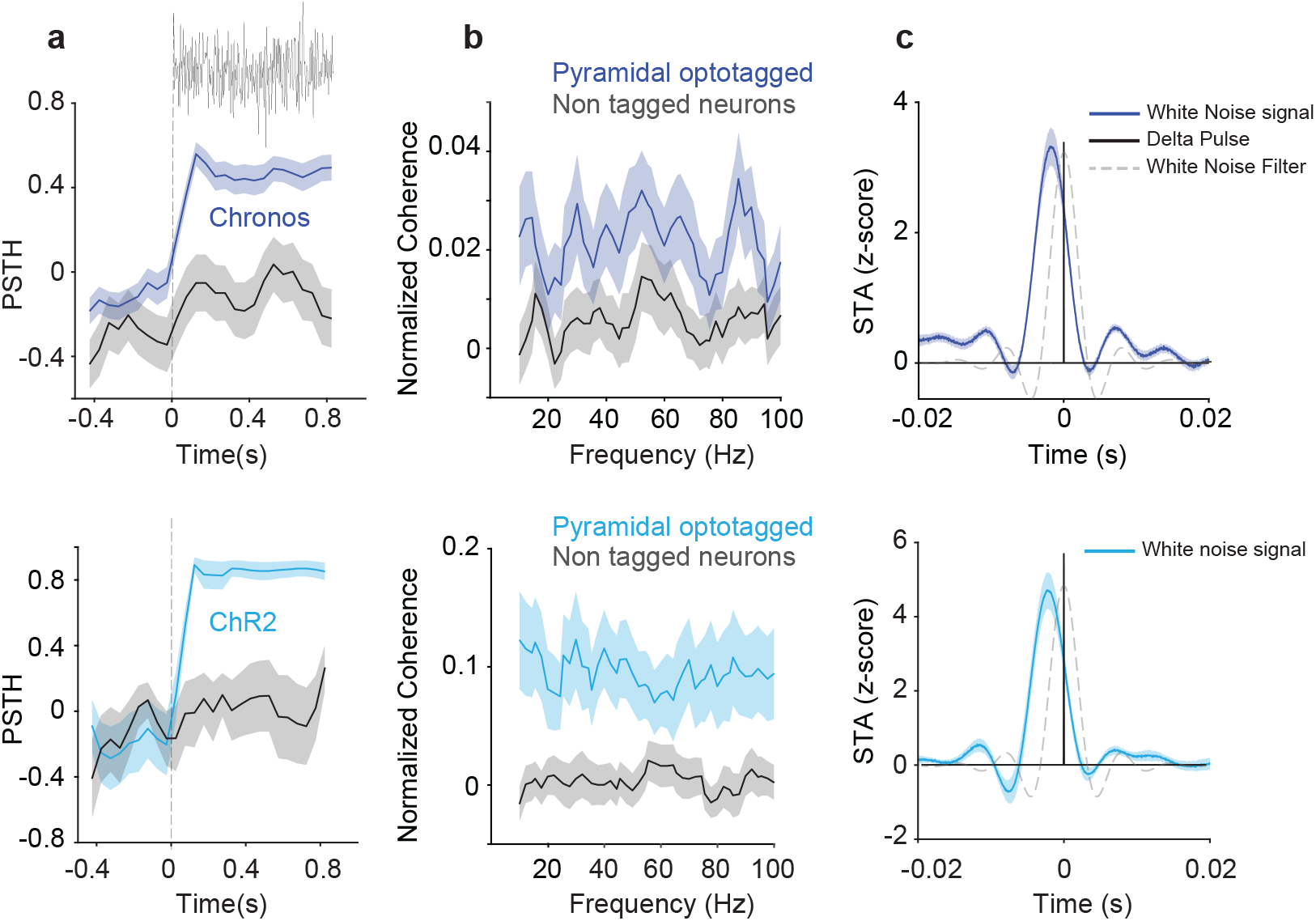
Sustained neuronal responses and absence of entrainment when cells (expressing Chronos or ChR2) were stimulated with white noise laser stimuli. **a**) White noise stimuli evoked a clear sustained response (similar to what was observed with DC stimuli) when pyramidal cells expressed Chronos or Chr2. **b**) The coherence spectrum between spike trains and the laser stimulus was flat for pyramidal optotagged cells expressing Chronos or Ch2. As a control, we show the coherence spectrum for the non-optotagged neurons, which shows much weaker coherence. **c**) Spike-triggered average of the white-noise stimulus around each spike, which shows that spikes are preceded by rapid depolarizations in the white noise occurring a few ms before. Note that the short-lived rhythmic modulation at a high frequency is comparable to the filter kernel used for the white-noise. Thus, the spike-triggered average for the white noise stimuli did not present additional oscillatory modulation, in agreement with the absence of gamma resonance when stimulating pyramidal cells optogenetically using Chronos or Chr2.

## Notes

### Competing Interest Statement

The authors have declared no competing interest.

## References

Boyden, E.S., Zhang, F., Bamberg, E., Nagel, G., Deisseroth, K., 2005. Millisecond-timescale, genetically targeted optical control of neural activity., 1263–1268 doi:10.1038/nn1525. ISSN: 1097-6256 Issue: 9 Journal Abbreviation: Nat Neurosci Publication Title: Nature neuroscience Volume: 8.

Bryant, H.L., Segundo, J.P., 1976. Spike initiation by transmembrane current: a white-noise analysis. 260, 279–314. URL:https://onlinelibrary.wiley.com/doi/abs/10.1113/jphysiol.1976.sp011516, doi: 10.1113/jphysiol.1976.sp011516._eprint: https://onlinelibrary.wiley.com/doi/pdf/10.1113/jphysiol.1976.sp011516.

Buzsáki, G., Draguhn, A., 2004. Neuronal oscillations in cortical networks. Science 304, 1926–1929.

Buzsaki, G., Voroslakos, M., 2023. Brain rhythms have come of age. Neuron.

Buzsáki, G., Wang, X.J., 2012. Mechanisms of Gamma Oscillations. Ann. Rev. Neurosci. 35, 203–225. doi:10.1146/annurev-neuro-062111-150444.

Cardin, J.A., Carlén, M., Meletis, K., Knoblich, U., Zhang, F., Deisseroth, K., Tsai, L.H., Moore, C.I., 2009. Driving fast-spiking cells induces gamma rhythm and controls sensory responses. Nature 459, 663–7. doi: 10.1038/nature08002.

Cardin, J.A., Palmer, L.A., Contreras, D., 2005. Stimulus-dependent gamma (30-50 Hz) oscillations in simple and complex fast rhythmic bursting cells in primary visual cortex. J Neurosci 25, 5339–50. doi: 10.1523/JNEUROSCI.0374-05.2005.

Charles, A.S., Park, M., Weller, J.P., Horwitz, G.D., Pillow, J.W., 2018. Dethroning the fano factor: A flexible, model-based approach to partitioning neural variability 30, 1012–1045. URL:https://doi.org/10.1162/neco_a_01062, doi: 10.1162/neco_a_01062.

Eden, U.T., Kramer, M.A., 2010. Drawing inferences from fano factor calculations 190, 149–152. URL:https://www.sciencedirect.com/science/article/pii/S0165027010001986, doi: 10.1016/j.jneumeth.2010.04.012.

Emiliani, V., Entcheva, E., Hedrich, R., Hegemann, P., Konrad, K.R., Lüscher, C., Mahn, M., Pan, Z.H., Sims, R.R., Vierock, J., Yizhar, O., 2022. Optogenetics for light control of biological systems 2, 1–25. URL: https://www.nature.com/articles/s43586-022-00136-4, doi: 10.1038/s43586-022-S0136-4. number: 1 Publisher: Nature Publishing Group.

Fries, P., 2015. Rhythm for Cognition: Communication Through Coherence. Neuron 88, 220–235. doi: 10.1016/j.neuron.2015.09.034.Rhythms,arXiv:15334406.

Gray, C., McCormick, D., 1996. Chattering cells: superficial pyramidal neurons contributing to the generation of synchronous oscillations in the visual cortex. Science 274, 109.

Grossberger, L., Battaglia, F.P., Vinck, M., 2018. Unsupervised clustering of temporal patterns in high-dimensional neuronal ensembles using a novel dissimilarity measure 14, e1006283. URL: https://journals.plos.org/ploscompbiol/article?id=10.1371/journal.pcbi.1006283, doi: 10.1371/journal.pcbi.1006283. publisher: Public Library of Science.

Hass, C.A., Glickfeld, L.L., 2016. High-fidelity optical excitation of cortico-cortical projections at physiological frequencies 116, 2056–2066. doi: 10.1152/jn.00456.2016.

Hutcheon, B., Yarom, Y., 2000. Resonance, oscillation and the intrinsic frequency preferences of neurons. Trends in neurosciences 23, 216–222.

Izhikevich, E.M., Desai, N.S., Walcott, E.C., Hoppensteadt, F.C., 2003. Bursts as a unit of neural information: selective communication via resonance. Trends in neurosciences 26, 161–167.

Jun, N.Y., Cardin, J.A., 2020. Activation of distinct channelrhodopsin variants engages different patterns of network activity. ENeuro 7.

Klapoetke, N.C., Murata, Y., Kim, S.S., Pulver, S.R., Birdsey-Benson, A., Cho, Y.K., Morimoto, T.K., Chuong, A.S., Carpenter, E.J., Tian, Z., et al., 2014. Independent optical excitation of distinct neural populations. Nature methods 11, 338–346.

Lewis, C.M., Ni, J., Wunderle, T., Jendritza, P., Lazar, A., Diester, I., Fries, P., 2021. Cortical gamma-band resonance preferentially transmits coherent input. Cell reports 35, 109083.

Mainen, Z.F., Sejnowski, T.J., 1995. Reliability of spike timing in neocortical neurons. Science 268, 1503–1506.

McGinley, M.J., Vinck, M., Reimer, J., Batista-Brito, R., Zagha, E., Cadwell, C.R., Tolias, A.S., Cardin, J.A., McCormick, D.A., 2015. Waking State: Rapid Variations Modulate Neural and Behavioral Responses. Neuron 87, 1143–1161. doi: 10.1016/j.neuron.2015.09.012.

Ni, J., Lewis, C., Wunderle, T., Jendritza, P., Diester, I., Fries, P., 2017. Gamma-band resonance of visual cortex to optogenetic stimulation. BioRxiv, 135467.

Niell, C.M., Stryker, M.P., 2010. Modulation of visual responses by behavioral state in mouse visual cortex. Neuron 65, 472–479.

Onorato, I., Neuenschwander, S., Hoy, J., Lima, B., Rocha, K.S., Broggini, A.C., Uran, C., Spyropoulos, G., Klon-Lipok, J., Womelsdorf, T., Fries, P., Niell, C., Singer, W., Vinck, M., 2020. A distinct class of bursting neurons with strong gamma synchronization and stimulus selectivity in monkey V1. Neuron 105, 180–197.

Pike, F., Goddard, R., Suckling, J., Ganter, P., Kasthuri, N., Paulsen, O., 2000. Distinct frequency preferences of different types of rat hippocampal neurones in response to oscillatory input currents. The Journal of Physiology 529, 205.

Rajdl, K., Lansky, P., Kostal, L., 2020. Fano factor: A potentially useful information 14. URL:https://www.frontiersin.org/articles/10.3389/fncom.2020.569049.

Rusch, T.K., Mishra, S., 2020. Coupled oscillatory recurrent neural network (cornn): An accurate and (gradient) stable architecture for learning long time dependencies. arXiv preprint arXiv:2010.00951.

Sirota, A., Montgomery, S., Fujisawa, S., Isomura, Y., Zugaro, M., Buzsáki, G., 2008. Entrainment of neocortical neurons and gamma oscillations by the hippocampal theta rhythm. Neuron 60, 683–697.

Sotomayor-Gómez, B., Battaglia, F.P., Vinck, M., 2023. Spike Ship: A method for fast, unsupervised discovery of high-dimensional neural spiking patterns URL:https://www.biorxiv.org/content/10.1101/2020.06.03.131573v4, doi: 10.1101/2020.06.03.131573. pages: 2020.06.03.131573 Section: New Results.

Spyropoulos, G., Saponati, M., Dowdall, J.R., Schölvinck, M.L., Bosman, C.A., Lima, B., Peter, A., Onorato, I., Klon-Lipok, J., Roese, R., et al., 2022. Spontaneous variability in gamma dynamics described by a damped harmonic oscillator driven by noise. Nature communications 13, 1–18.

Stark, E., Eichler, R., Roux, L., Fujisawa, S., Rotstein, H.G., Buzsáki, G., 2013. Inhibition-induced theta resonance in cortical circuits. Neuron 80, 1263–1276.

Steinmetz, N.A., Aydin, C., Lebedeva, A., Okun, M., Pachitariu, M., Bauza, M., Beau, M., Bhagat, J., Böhm, C., Broux, M., Chen, S., Colonell, J., Gardner, R.J., Karsh, B., Kloosterman, F., Kostadinov, D., Mora-Lopez, C., O’Callaghan, J., Park, J., Putzeys, J., Sauerbrei, B., van Daal, R.J.J., Vollan, A.Z., Wang, S., Welkenhuysen, M., Ye, Z., Dudman, J.T., Dutta, B., Hantman, A.W., Harris, K.D., Lee, A.K., Moser, E.I., O’Keefe, J., Renart, A., Svoboda, K., Häusser, M., Haesler, S., Carandini, M., Harris, T.D., 2021. Neuropixels 2.0: A miniaturized high-density probe for stable, long-term brain recordings. Science 372. doi: 10.1126/science.abf4588.

Vaidya, S.P., Johnston, D., 2013. Temporal synchrony and gamma-to-theta power conversion in the dendrites of ca1 pyramidal neurons. Nature neuroscience 16, 1812–1820.

Victor, J.D., Purpura, K.P., 1996. Nature and precision of temporal coding in visual cortex: a metric-space analysis 76, 1310–1326. doi: 10.1152/jn.1996.76.2.1310.

Victor, J.D., Purpura, K.P., 1997. Metric-space analysis of spike trains: theory, algorithms and application 8, 127–164. URL:https://doi.org/10.1088/0954-898X_8_2_003, doi: 10.1088/0954-898X_8_2_003. publisher: Taylor & Francis _eprint: https://doi.org/10.1088/0954-898X-8_2_003.

Vinck, M., Batista-Brito, R., Knoblich, U., Cardin, J.A., 2015. Arousal and Locomotion Make Distinct Contributions to Cortical Activity Patterns and Visual Encoding. Neuron 86, 740–754. doi:10.1016/j.neuron.2015.03.028.

Vinck, M., Battaglia, F.P., Womelsdorf, T., Pennartz, C., 2012. Improved measures of phase-coupling between spikes and the Local Field Potential. J Comput Neurosci 33, 53–75. doi:10.1007/s10827-011-0374-4.

Vinck, M., Uran, C., Spyropoulos, G., Onorato, I., Broggini, A., Schneider, M., Canales-Johnson, A., 2023. Principles of large-scale neural interactions. Neuron.

Voloh, B., Valiante, T.A., Everling, S., Womelsdorf, T., 2015. Theta-gamma coordination between anterior cingulate and prefrontal cortex indexes correct attention shifts. Proc. Natl. Acad. Sci. U. S. A. 112, 8457–8462.

Wang, Q., Gao, E., Burkhalter, A., 2011. Gateways of ventral and dorsal streams in mouse visual cortex 31, 1905–1918. URL:https://www.JNEUROSCI.org/content/31/5/1905, doi: 10.1523/JNEUROSCI.3488-10.2011. publisher: Society for Neuroscience Section: Articles.

Wang, W.T., Wan, Y.H., Zhu, J.L., Lei, G.S., Wang, Y.Y., Zhang, P., Hu, S.J., 2006. Theta-frequency membrane resonance and its ionic mechanisms in rat subicular pyramidal neurons 140, 45–55. doi: 10.1016/j.neuroscience.2006.01.033.

Wang, X.J., 2010. Neurophysiological and computational principles of cortical rhythms in cognition. Physiol. Rev. 90, 1195–1268.

Yu, C., Cassar, I.R., Sambangi, J., Grill, W.M., 2020. Frequency-specific optogenetic deep brain stimulation of subthalamic nucleus improves parkinsonian motor behaviors 40, 4323. URL:http://www.jneurosci.org/content/40/22/4323.abstract, doi: 10.1523/JNEUROSCI.3071-19.2020.

